# Speciation trajectories in recombining bacterial species

**DOI:** 10.1101/095885

**Authors:** Pekka Marttinen, William P. Hanage

## Abstract

While speciation in eukaryotes is well-studied, a controversy over the nature of bacterial species due to recombination between distant strains continues. It is generally agreed that bacterial diversity can be classified into genetically and ecologically cohesive units, but what produces such variation is a topic of intensive research. Recombination may maintain coherent species of frequently recombining bacteria, but the emergence of distinct clusters within a recombining species, and the impact of habitat structure in this process are not well described, limiting our understanding of how new species are created. Here we present a model of bacterial evolution in overlapping habitat space. We show three different outcomes are possible for a pair of clusters, depending on the size of their habitat overlap: fast divergence with little interaction between the clusters, slow divergence with frequent recombination between the clusters, and stationary or near stationary population structure, where the clusters remain at an equilibrium distance for an indefinite time. We fit our model to two data sets. In Streptococcus pneumoniae, we find a genomically and ecologically distinct subset, held at a relatively constant genetic distance from the majority of the population through frequent recombination with it, while in Campylobacter jejuni, we find a minority population we predict will continue to diverge at a higher rate. This approach may predict and define speciation trajectories in multiple bacterial species.

---

While speciation in eukaryotes is well-studied (Coyne *et al.*, 2004), a controversy over the nature of bacterial species due to recombination between distant strains continues (Fraser *et al.*, 2007). It is generally agreed that bacterial diversity can be classified into genetically and ecologically cohesive units (Vos, 2011; Caro-Quintero and Konstantinidis, 2012; Shapiro and Polz, 2014), but what produces such variation is a topic of intensive research (Cohan and Perry, 2007; Shapiro, 2014; Shapiro *et al.*, 2016). Recombination may maintain coherent species of frequently recombining bacteria (Fraser *et al.*, 2009; Marttinen *et al.*, 2015; Dixit *et al.*, 2016), but the emergence of distinct clusters within a recombining species, and the impact of habitat structure in this process are not well described, limiting our understanding of how new species are created. Here we present a model of bacterial evolution in overlapping habitat space. We show three different outcomes are possible for a pair of clusters, depending on the size of their habitat overlap: fast divergence with little interaction between the clusters, slow divergence with frequent recombination between the clusters, and stationary or near stationary population structure, where the clusters remain at an equilibrium distance for an indefinite time. We fit our model to two data sets. In *Streptococcus pneumoniae*, we find a genomically and ecologically distinct subset, held at a relatively constant genetic distance from the majority of the population through frequent recombination with it, while in *Campylobacter jejuni*, we find a minority population we predict will continue to diverge at a higher rate. This approach may predict and define speciation trajectories in multiple bacterial species.

Several explanations have been offered for the genetic and ecological differentiation between bacterial populations. In the Ecotype Model (Cohan and Perry, 2007), ecological niche-specific adaptive mutations cause genome-wide selective sweeps that purge variability among isolates in the same the niche, resulting in genetically differentiated clusters that correspond to niches. While recombination may maintain genetic coherence, as discussed above, theory suggests selection is necessary for genetic diversification (Polz *et al.*, 2013). Recently, a model of ecological differentiation among sympatric recombining bacteria has been developed (Shapiro *et al.*, 2012; Friedman *et al.*, 2013). The model assumes an initial freely recombining population, and the differentiation is triggered by an acquisition of a few habitat-specific alleles through horizontal gene transfer. If the habitat-specificity results in varying recombination rates within and between habitats, for example due to increased physical proximity or sequence similarity, the result is gradual diversification across the genome, eventually creating genomically and ecologically distinct clusters. Unlike in the Ecotype Model, which assumes genome-wide sweeps, here the sweeps occur only for the habitat-specific genes, but the overall genetic differentiation does not happen immediately, thanks to frequent recombination that breaks the linkage between the habitat specific genes and the rest of the genome. The resulting pattern has a small number of short genome regions with strong habitat association, while the majority of the genome shows little correlation with the habitat, and such a pattern was observed between a pair of closely-related *Vibrio* bacteria (Shapiro *et al.*, 2012).

Fig. 1 shows population structures in data sets with 616 *Streptococcus pneumoniae* (Croucher *et al.*, 2013) and 235 *Campylobacter jejuni* samples (Sheppard *et al.*, 2013, 2014; Cody *et al.*, 2013) (see Methods). Both include strains divergent from the rest of the population, providing us with an opportunity to investigate the early stages of bacterial differentiation. In particular, the *S. pneumoniae* data consists of 16 sequence clusters (SCs) of which one, SC12, differs from the rest, and has previously been characterized as atypical pneumococci representing a distinct species (Croucher *et al.*, 2013, 2014). All other SCs are at the same equilibrium distance from each other, maintained by recombination, corresponding to the main mode in the distance distribution (Marttinen *et al.*, 2015). Two additional modes can be discerned: one close to the origin comprising the within SC distances, which may be explained by selection of some sort, and the other representing the broad division of the data into SC12 vs. rest, which indicates less frequent recombination between these two clusters. Whether SC12 is a nascent cluster, which will continue to diverge, is not known. It is also possible that the distance could be an equilibrium produced by the combination of mutational divergence and occasional recombination with the parent cluster. A similar minor mode is found in *C. jejuni*, in this case arising from a single divergent isolate shown in red. Whether this is an isolate from a cluster in the early stages of divergence is similarly unknown.

**Figure 1:**
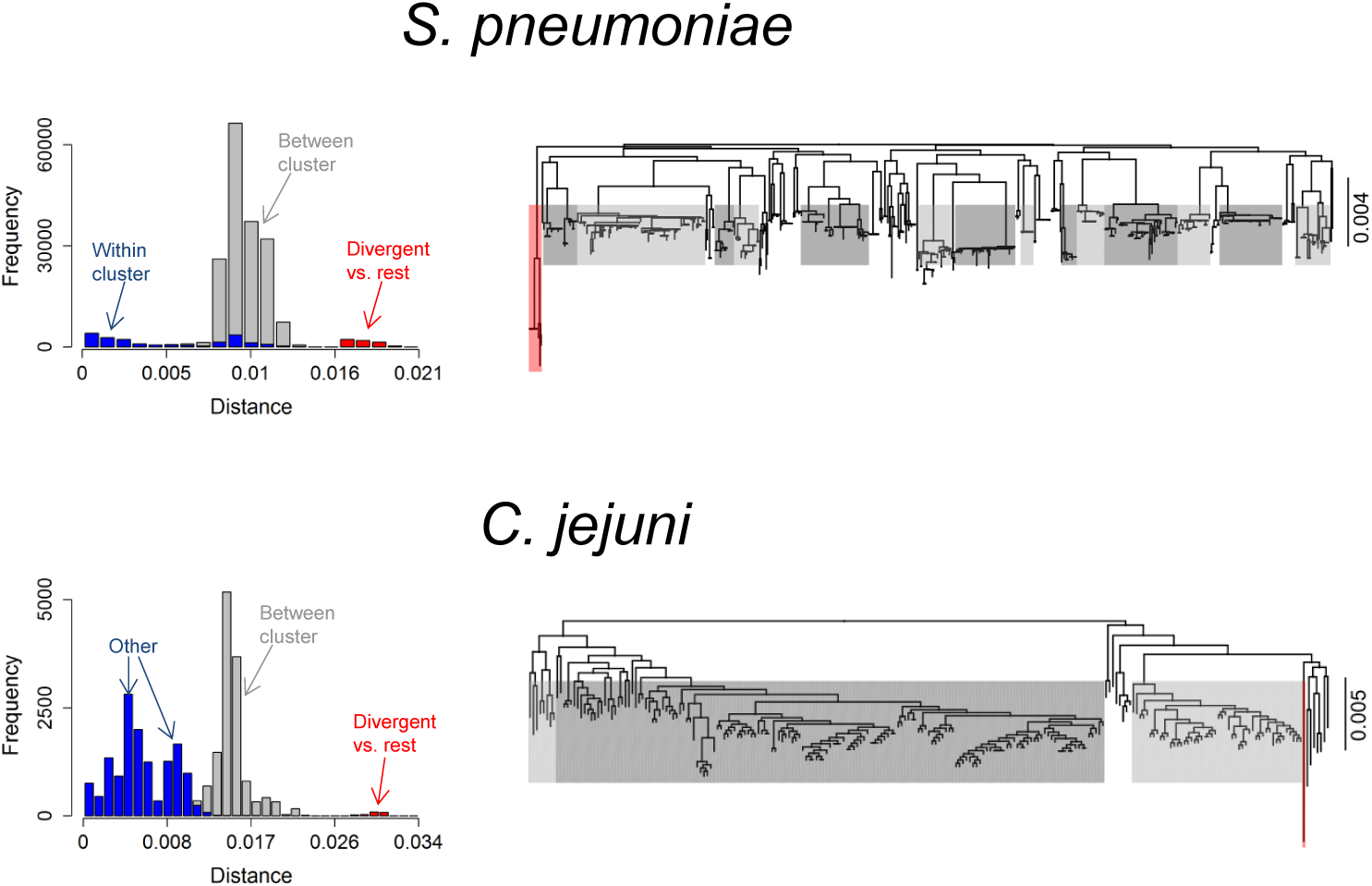
Population structures in *S. pneumoniae* and *C. jejuni* data sets. The left panels show distributions of pairwise distances computed between all strain pairs in the data sets, and the right panels show the phylogenies. In the *S. pneumoniae* phylogeny, previously identified 16 sequence clusters are annotated as follows: the divergent cluster with red, 14 other monophyletic clusters with gray, and the remaining non-monophyletic cluster is not colored. Distances within and between these clusters are annotated in the distance histogram. Similarly, for *C. jejuni*, three clusters corresponding to separate branches of the phylogeny are colored with gray and one divergent strain with red, and the distances within and between these clusters are shown in the histogram. Annotation “Other” refers to within cluster comparisons as well as distances between the non-colored strains and other strains.

The goal to understand the population sub-divisions observed in Fig. 1 motivated us to develop a model that could re-produce similar patterns. Previously, models have been used to investigate the impact of homologous recombination on population structure (Fraser *et al.*, 2009; Doroghazi and Buckley, 2011), the distribution of accessory genome (Baumdicker *et al.*, 2012; Lobkovsky *et al.*, 2013; Collins and Higgs, 2012), parallel evolution of the core and accessory genomes (Marttinen *et al.*, 2015), and the spread of antibiotic resistance (Niehus *et al.*, 2015). Here we extend the model of sympatric differentiation (Shapiro *et al.*, 2012; Friedman *et al.*, 2013) in two ways. First, we introduce an explicit, controllable barrier for recombination between the two populations, and second, we derive an analytical approximation of the model. An outline of the resulting ‘Overlapping Habitats Model’, is shown in Fig. 2. Its key characteristic is the existence of two populations of different types of strains living in partially overlapping habitats. Recombination between the populations only occurs between individuals in the shared habitat, while migration enables strains to move between strain type specific and shared parts of the habitat space (see Methods). The explicit barrier for recombination together with the analytical approximation make it for the first time possible to do inference about the amount of interaction between the populations. In particular, the model allows us to rapidly predict how the population structure will evolve given a certain amount of habitat overlap, and, on the other hand, to learn the parameters resulting in a given equilibrium population structure.

**Figure 2:**
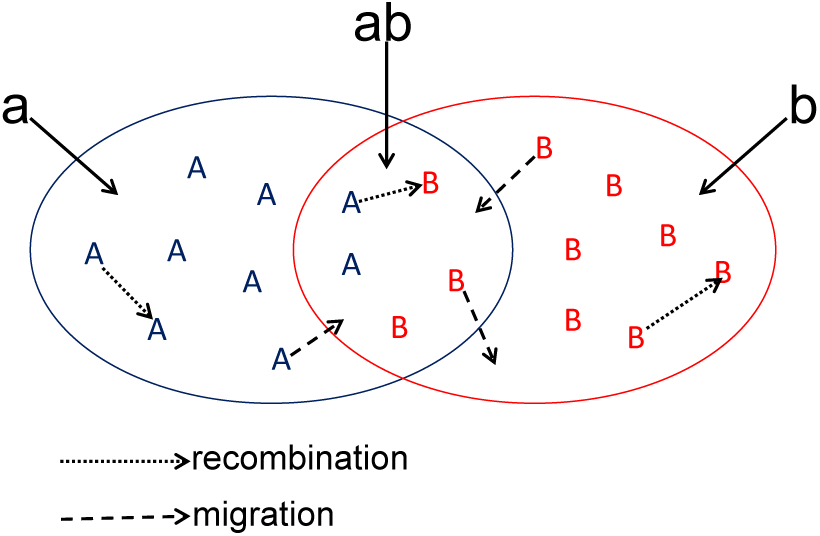
Outline of the Overlapping Habitats Model. The model assumes two types of strains, *A* and *B*, that live in habitats *a* or *b*, respectively. In addition, both types can live in the intersection of the habitats, denoted as *ab*. Type *A* strains can migrate between *a* and *ab* and type *B* strains between *b* and *ab*. Strain can recombine with other strains in the same habitat.

To investigate the impact of habitat structure on population structure, we simulated the Overlapping Habitats Model for 100,000 generations with two clusters, each with 5,000 strains. We varied the amount of habitat overlap and migration, but used realistic mutation and recombination rates (see Methods). Fig. 3 shows how the within and between cluster distances evolved during the simulation. As expected with the smallest overlap (left-most panels), the limited interaction resulted in rapid divergence of the clusters, although within cluster distances reached an equilibrium as expected (Fraser *et al.*, 2007; Marttinen *et al.*, 2015). With the largest overlap (right panels) two distinguishable clusters emerged, with the between cluster distance exceeding the within distance. However the clusters did not proceed to full separation, but rather maintained an equilibrium level of separation for what appeared to be an indefinitely long time. With an intermediate overlap the simulation still had characteristics of the stationary behaviour; however, now the clusters slowly drifted apart as a result of genes one by one escaping the equilibrium. To understand the equilibrium, we first note that if the clusters are already very close, then a recombination event between them does not make them any more similar. If the clusters are very distant, the ability to recombine vanishes. Intuitively, the equilibrium, if such exists, is located at an intermediate distance where the cohesive force of recombination equals the diversifying force of mutation.

**Figure 3:**
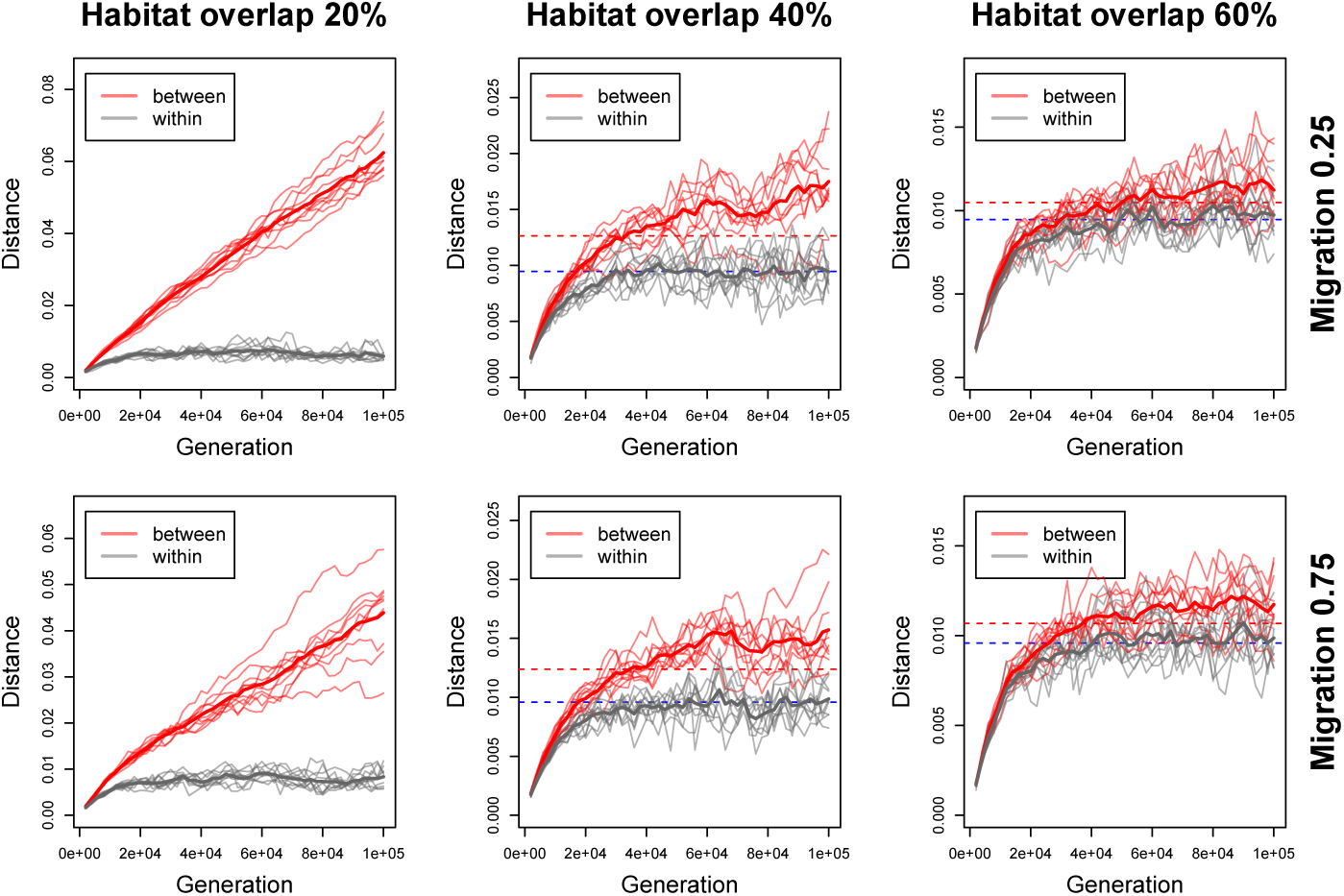
Simulation results from the Overlapping Habitats Model. The figure shows the evolution of distances within and between strain types in simulations with 10^6^ generations. The solid thin red and gray lines show the median between and within strain type distances in ten repetitions, and the thick lines show the averages across the repetitions. The dashed horizontal lines show the predicted equilibrium distances from the deterministic approximation. If no dotted line is shown, the deterministic model did not have a solution.

We next investigated which of the three alternative types of differentiation: equilibrium, slow divergence, or rapid clonal divergence, best explains the population divisions in the *S. pneumoniae* and *C. jejuni* data (Fig. 1). To fit the Overlapping Habitats Model, which represents the equilibrium or slow divergence cases, we tentatively assumed the distances between the divergent strains and other strains to be at equilibrium, and used a plug-in recombination rate estimate from the literature to compute the approximate overlap that would produce the observed level of separation (see Methods). For both data sets, a simulation with these parameters resulted in two separate clusters that were diverging slowly, with rates of 0.32 (*S. pneumoniae*) and 0.45 (*C. jejuni*) relative to the clonal divergence rates. This indicates the separation between the clusters, especially in the *C. jejuni* data, has exceeded the level where recombination could prevent the divergence, and, consequently, the equilibrium distance is easy to escape. However, these results alone do not yet allow us to separate the two possible explanations: first, the clusters are in the process of slow divergence, as just described, or second, the clusters are in the process of rapid clonal diversification, and the distance between them just happens momentarily to be as observed.

A detailed comparison of the models’ outputs revealed a systematic difference in the ecoSNP distributions between the scenarios of clonal divergence vs. equilibrium or slow divergence, where ecoSNPs are defined, as in (Shapiro *et al.*, 2012), as variants present in all strains of one cluster and absent from all strains of the other cluster. In particular, with rapid divergence and little recombination between the clusters, the ecoSNPs were rather uniformly distributed across the genes (Fig. 4). On the other hand, under the equilibrium the majority of ecoSNPs were concentrated in only a few genes that already had escaped the equilibrium, while the majority of genes had no ecoSNPs at all. For both data sets, the ecoSNP distribution supports the interpretation that the observed population structure is a result of equilibrium or slow divergence, rather than rapid clonal divergence. In the *S. pneumoniae* data the observed proportion of genes with no ecoSNPs is even higher than predicted by the overlap model, suggesting that previously published recombination rates may be underestimates.

**Figure 4:**
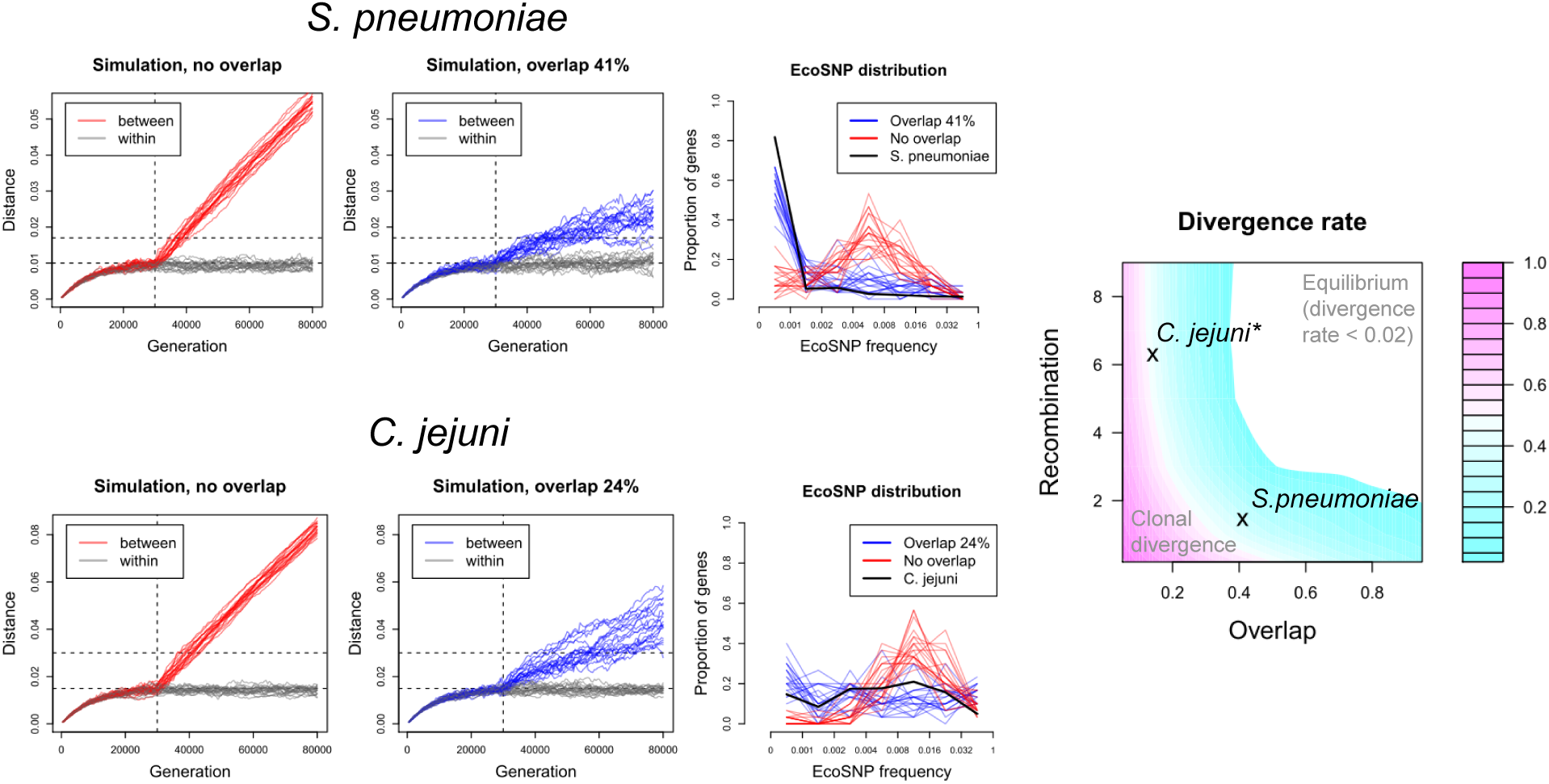
Comparing model output with the *S. pneumoniae* and *C. jejuni* data, and a summary of divergence rates. For each data set, we simulated the Overlapping Habitats Model 20 times without overlap (leftmost small panels) and with the estimated overlap (center small panels). A barrier representing the size of the overlap between the clusters was introduced at the 30,000th generation (dashed vertical line) after which the clusters diverged. The horizontal lines show for reference the within and between cluster distances in *S. pneumoniae* and *C. jejuni*. Simulated ecoSNP distributions with and without overlap, computed at the generation when the simulated between-cluster distance matched the observed value, are compared with the observed ecoSNP distributions (rightmost small panels). The panel on the right summarizes the simulated rate of divergence between the two clusters relative to the clonal divergence. (*the heatmap is based on the mutation rate in *S. pneumoniae*, and, therefore, the location of *C. jejuni* is modified by moving it to the closest contour line corresponding to the divergence rate estimated using the correct mutation rate, for which results are shown in Supplementary Fig. S3)

We note that the concentration of ecoSNPs in a few genome regions has previously been taken as evidence for gene-specific sweeps of habitat-specific adaptive alleles acquired through horizontal gene transfer (Shapiro *et al.*, 2012). Our results suggest a similar pattern may emerge as a result of drift making a region divergent enough between the clusters to reduce their ability to recombine within the region, after which the region continues rapid diversification while the rest of the genome remains at equilibrium. This recalls the concept of ‘fragmented’ speciation in which different parts of the genome speciate at different times (Retchless and Lawrence, 2010), except in this case this can be achieved without explicit selective processes on the diverging region. Eventually this results in highly divergent habitat-specific loci surrounded by regions with little habitat association. In practice this process could happen together with selection at the habitat-specific loci, as both processes have the potential to increase differentiation and create ecoSNPs between the clusters. We note that while quantitatively the simulation output obviously depends on the exact parameter values, qualitatively the conclusions regarding the main patterns observed in the data sets seem robust across a wide range of parameter values.

We have introduced the Overlapping Habitats Model and shown that with realistic parameter values stationary or nearly stationary population divisions may emerge, creating what might be termed ‘satellite species’ as seen in *S. pneumoniae*, and that these may be distinguished from dynamically diverging clusters using ecoSNPs, as shown by the analysis of *C. jejuni*. In our model the habitat could represent, for example, physical separatedness, biochemical properties, or any abstract division of the space according to the Hutchinson’s *n*-dimensional niche (Hutchinson, 1957). Our model is mainly about recombination; the ecological and selective aspects are implicit and follow from the division of the habitat-space into regions suitable for different strain types. The habitat-specificity is assumed heritable and non-mutable, and could in practice be caused by a small number of genes. Despite the simplicity of the model, it adequately captured the main sub-divisions in two data sets. Nonetheless, much of the ecological and genomic structure in the data will not be captured by the model, for example the individual sequence clusters within the main group in the *S. pneumoniae* data. Our model does not contradict with this additional structure, but instead shows that the individual sequence clusters can indeed be ecologically different, and still maintain the equilibrium distance between them, as a mere 60% habitat overlap already is sufficient for this (Fig. 3). Our model provides means to characterize equilibrium structures in bacterial populations and we believe it will be helpful to understand similar patterns in many other bacterial genomic data sets.

## Methods

### Data

Core gene alignments and the cluster annotation of the *S. pneumoniae* strains were obtained from (Croucher *et al.*, 2013). As an additional data cleaning step, we removed all genes whose alignment length was less than 265bp, which corresponded to the 0.05th quantile of the lengths of the alignments of the core genes. This step was added to increase confidence in the genes detected. This left us with 1,191 core genes in the 616 pneumococcal isolates. More specifically, the genes are here clusters of orthologous genes (COGs), and we use these terms interchangeably.

The *C. jejuni* data consisted of 239 previously published genomes (Sheppard *et al.*, 2013, 2014; Cody *et al.*, 2013). From the reference-based assemblies mapped to the NCTC11168 reference genome, we extracted 423 COGs using ROARY (Page *et al.*, 2015) with default settings. As a data cleaning step, we removed four isolates with significantly increased levels of missing data. Additionally, we removed COGs whose alignment lengths were less than the 0.05th quantile (225bp) of all lengths. This left us with 401 COGs with 235 isolates. The divergent isolate in Fig. 1 differs from others in terms of its sampling location (New Zealand), and by being the only isolate sampled from ‘environment’ and having ST=2381.

### Simulation model

As the basis of our model, we use a Wright-Fisher forward simulation of discrete generations, where each generation is sampled with replacement from strains in the previous generation. In our model, a strain is represented by a collection of genes, similar to (Fraser *et al.*, 2007), and we assume the genes are ‘core’, i.e., present in all strains. Each gene is encoded as a binary sequence of fixed length (500 bp). Our model has in total four free parameters: mutation rate, homologous recombination rate, the proportion of habitat overlap, and migration rate. Mutations and recombinations are assumed to take place between sampling of the generations. Mutations change one base in the target sequence, while recombination is assumed to result in the whole gene of the recipient to be replaced by the corresponding gene of the donor. Recombination is allowed only between strains within the same habitat, and accepted with probability that declines with respect to increasing sequence divergence (Zawadzki *et al.*, 1995; Vulić *et al.*, 1997; Majewski *et al.*, 2000). As opposed to (Fraser *et al.*, 2007; Marttinen *et al.*, 2015), we simulate complete binary sequences, avoiding the need for additional approximations.

A population of strains of two types, *A* and *B*, is simulated. We assume the strain types live in different habitats, such that type *A* strains live in habitat *a* and type *B* strains in habitat *b*; however, part of the habitat space is shared such that both strain types can inhabit it, and we denote the shared part by *ab*. Habitat-specificity encoding genes are assumed implicit and not simulated in our model, and we assume that a strain type can not be changed by recombination or mutation. Migration of type *A* strains between habitats *a* and *ab* is achieved by sampling the next generation of strains in *a*, for example, from all type A strains such that strains in *ab* are sampled with a relative weight determined by the migration parameter. This corresponds to the assumption that strains within each habitat compete against each other and those trying to enter the habitat. In detail, the sampling scheme can be described as follows. We denote by *A_a_* and *A_ab_* type A strains that are currently in *a* or *ab* environments; *B_b_* and *B_ab_* are defined correspondingly. We sample strains for *a* with replacement from *A_a_* and *A_ab_* such that the probability of sampling a strain *x* is equal to

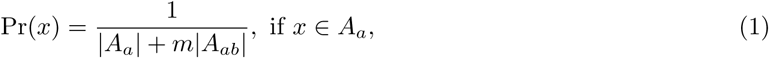

and

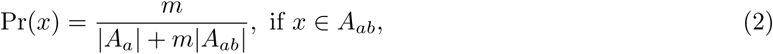

where 0 ≤ *m* ≤ 1 is the migration parameter. Value *m* = 0 corresponds to no migration, in which case Equations 1 and 2 reduce to sampling the next generation for environment *a* from strains already in that environment. On the other hand, *m* =1 corresponds to unlimited migration, and the next generation is sampled with equal probability from all type *A* strains in both environments *a* and *ab*. Strains for the *b* environment are sampled similarly from strains in *b* and *ab* environments. Finally, strains for the *ab* environment are sampled according to

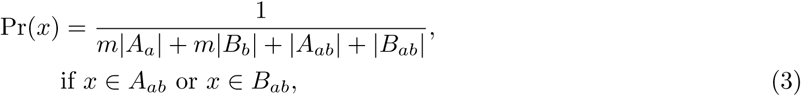

and

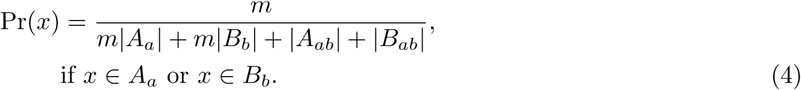

Thus, if *m* = 0, the next generation of strains for the *ab* environment is sampled from strains already in the environment. In the other extreme (*m* = 1), the strains are sampled from all strains in both populations.

### Deterministic approximation of the model

We also derive a deterministic approximation of the Overlapping Habitats Model, which enables rapid prediction of the evolution of the population structure without simulating the actual sequences. The model is based on average distances between and within the different sub-groups of the whole population: *A_a_*, *A_ab_*, *B_ab_*, and *B_b_* (see the previous sub-section). In detail, let **d** be a vector comprising all 4 within and 6 between distances possible for the four groups. In the Supplementary Text S1, we derive a function *f*, that expresses how the average distances in the next generation, **d**\*, approximately depend on the distances **d** in the current generation:

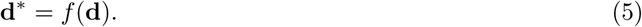

One of the main interests is to identify stationary points in the distance distribution, i.e., distances **d**, for which

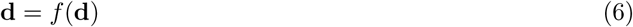

holds.

We have implemented two methods to solve Equation (6). The first consists of using the update rule (5) repeatedly until the **d** converges, in which case the stationarity condition (6) is satisfied. The second way to solve (6) is to use a quasi-Newton method, implemented in the *optim*-function of the **R** software, to minimize the objective function *h*, defined as follows:

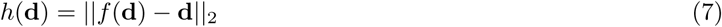

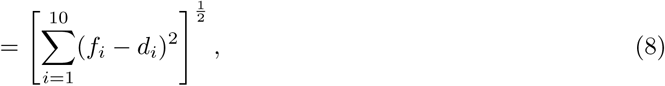

where *f_i_* is the prediction for the *i*th element in the distance vector of the next generation, and *d_i_* the current value of the corresponding element. In practice, we have reached the best performance by first running the Newton’s method, which is fast, followed by the robust sequential update procedure to confirm convergence.

Investigation of Fig. 3 reveals that the deterministic approximation predicts the within cluster distances observed in simulation with high accuracy. Also, with the smallest overlap, the deterministic approximation does not have a solution, allowing us to immediately predict the rapid divergence seen in the simulation. However, we also see that the approximation has a tendency to underestimate the between cluster distances compared to the simulation. The reason for this is that the deterministic approximation is based on average distances within and between the populations, and therefore it does not account for variation in distances between specific donor and recipient alleles, resulting in overestimation of the impact of recombination. For the same reason it is not possible to use the approximation to determine how easy it is to escape the equilibrium mode in the distance distribution. Therefore, we adopted in our analyses a strategy to first estimate the parameters with the deterministic approximation (see below), and then run the simulation model with the learned parameter values to produce the final detailed prediction.

### Model fitting

As discussed above, the *S. pneumoniae* data can be broadly divided into two sub-populations. To estimate the habitat overlap, we tentatively assumed the population structure, i.e., the within and between sub-population distances observed, represented an equilibrium. We fitted the model by solving the deterministic formula to determine parameter values that produced the distances in the data (*within*=0.01, *between*=0.017) as a stationary condition (Fig. S2). To determine the remaining parameters, we set the recombination rate, *r*/*m* to a previously reported value *r*/*m* = 11.3 (Croucher *et al.*, 2013). The proportion of diverging strains of the whole population was set to 5%, and migration to 0.5 (results were insensitive to these choices, see Fig. S2). These specifications led to an estimate of 41% habitat overlap.

The parameters for the *C. jejuni* were estimated similarly. In detail, we assumed that the *within* population distance was 0.015 (the main mode) in the data and the *between* distance 0.03 (the small separate mode). We fixed the recombination rate to a plug-in estimate of *r*/*m* = 49, derived from an estimate that 98 percent of substitutions in MLST genes in the species are due to recombinations (Yu *et al.*, 2012). We again set the proportion of the diverging strains to be 5% of the whole population. These specifications yielded an estimate of 24% habitat overlap.

For both data sets, we set the total number of strains simulated as 10,000 and the number of genes as 30. As each gene had length 500, this corresponded to the total genome size of 15,000 bp. The probability of accepting a recombination was assumed to decline log-linearly with respect to the distance between the alleles in the donor and recipient strains, according to 10^*−Ax*^, where *x* is the Hamming distance between the alleles. We used *A* = 18 for the parameter that determines the rate of the decline, according to empirical data, see Fraser *et al.* (2007).

## Code availability

R-code to run the model is available at https://users.ics.aalto.fi/~pemartti/habitat_simulation/.

## Acknowledgements

This work was funded by the Academy of Finland (grants no. 286607 and 294015 to PM). The authors thank Sam Sheppard, University of Oxford, for providing the *C. jejuni* data, and Brian Arnold, Harvard T.H. Chan School of Public Health, for assistance with data processing and helpful comments. The calculations presented above were performed using computer resources within the Aalto University School of Science “Science-IT” project.

